# A tunable microfluidic device enables cargo encapsulation by cell-or organelle-sized lipid vesicles comprising asymmetric lipid monolayers

**DOI:** 10.1101/534586

**Authors:** Valentin Romanov, John McCullough, Abhimanyu Sharma, Michael Vershinin, Bruce K. Gale, Adam Frost

**Author notes:** These authors contributed equally to this work (Valentin Romanov, John McCullough).

## Abstract

Cellular membranes play host to a wide variety of morphologically and chemically complex processes. Although model membranes, like liposomes, are already widely used to reconstitute and study these processes, better tools are needed for making model bilayers that faithfully mimic cellular membranes. Existing methods for fabricating cell-sized (μm) or organelle-sized (tens to hundreds of nm) lipid vesicles have distinctly different requirements. Of particular note for biology, it remains challenging for any technique to efficiently encapsulate fragile cargo molecules or to generate liposomes with stable, asymmetric lipid-leaflets within the bilayer. Here we describe a tunable microfluidic device and protocol for fabricating liposomes with desired diameters ranging from ~10 μm to ~100 nm. Lipid vesicle size is templated by the simple inclusion of a polycarbonate filter within the microfluidic system and tuned with flow rate. We show that the vesicles made with our device are stable, unilamellar, lipid asymmetric, and capable of supporting transmembrane protein assembly, peripheral membrane protein binding, as well as soluble cargo encapsulation (including designer nanocages for biotechnology applications). These fabricated vesicles provide a new platform for studying the biophysically rich processes found within lipid-lipid and lipid-protein systems typically associated with cellular membranes.

---

Filled with an aqueous solvent and bounded by a fluid lipid bilayer, liposomes are popular mimetics for studying biological membranes and membrane-associated biochemical activities. Giant Unilamellar Vesicles (GUVs, >1μm) are cell-sized liposomes. Among other processes, GUVs are routinely used for studying *in vitro* protein synthesis^[1]^, actin polymerization^[2]^ and curvature-dependent protein-lipid dynamics^[3]^. Large Unilamellar Vesicles (LUVs, < 1 μm) are organelle-sized liposomes. LUVs have also been used extensively to study the molecular structures and functions of membrane-binding proteins, including pore-forming toxins^[4]^, large GTPases of the dynamin family^[5]^, ESCRT protein complexes^[6]^, and BAR^[7]^ domain proteins among many others. Despite their utility, however, liposome manufacturing protocols and the lipid membrane properties that result from each method vary widely^[8]^. To address these shortcomings, we sought to develop a simple and tunable microfluidic device for generating unilamellar LUVs or GUVs with defined lumenal contents and asymmetric lipid-leaflet compositions. Such a device would enable researchers to reconstitute complex cellular phenomena *in vitro* for detailed characterizations.

Microfluidic techniques are effective for: 1) synthesizing liposomes of defined diameter ranges; 2) encapsulating a variety of solute molecules; 3) lowering reagent consumption, and 4) generating lipid bilayers that comprise mixtures of different lipid species. Final liposome size depends on the starting microfluidic droplet size, and this necessitates fabricating channels that are roughly the same size as the desired vesicles. However, fabrication of micro (> 1 μm) or nanoscale (< 1 μm) channels typically requires access to cleanroom facilities and fabrication expertise. Micro channels are also prone to clogging and require filtered samples and, in many cases, chemical functionalization of the channel to promote stable droplet formation. Fabrication and operation of smaller channels (< 1 μm) requires even finer control over device processing and fluidic control parameters. As a result, few techniques exist for the formation of LUVs and no technique currently exists that can be tuned to generate either GUVs or LUVs using the same methodology.

Here we report a microfluidic device and protocol for the fabrication of either cell-sized GUVs or organelle-sized LUVs. This microfluidic approach combines a *y*-mixer design with cross-flow filter emulsification to rapidly produce micro or nanoscale lipid-stabilized droplets that are subsequently converted to liposomes by the phase-transfer method^[9]^. Torque-balance modeling of droplet formation suggested that final liposome size depends on the polycarbonate filter pore size, allowing us to create ~ 8 μm liposomes (GUVs) using a 5 μm polycarbonate filter and ~ 300 nm liposomes (LUVs) using a 100 nm polycarbonate filter. Our experiments confirm that liposome size can be controlled by substituting polycarbonate filters with different pore sizes. By utilizing a number of chemical, biological and microscopy assays we show that at both scales the majority of generated liposomes are unilamellar and that only a small subset possess any solvent contamination as detected by electron cryo-microscopy. Furthermore, we verify the ability of LUVs created with a complex phospholipid composition to bind to, and be remodeled by, the ESCRT-III protein CHMP1B (Charged Multivesicular Body Protein 1B). Finally, we demonstrate that at either scale, liposomes made with our device can be used to encapsulate and protect both organic and synthetic macromolecular cargos under a variety of solution conditions.

## Results and Discussion

### Templating vesicle size

We tested whether embedding polycarbonate filters with defined pore sizes within a microfluidic device could be used to create liposomes at either the micro or nanoscale using the Phase-Transfer (PT) method^[9,10]^. In this approach, a dispersed phase (solution to be encapsulated) is driven through a rigid filter into a shearing continuous phase (immiscible with the dispersed phase) under cross-flow emulsification conditions leading to the formation of droplets (Figure 1a). Droplet size primarily depends on the membrane pore size and can be tuned by adjusting the wall shear stress (which is a function of cross-flow velocity and fluid properties)^[11]^. However, beyond a certain cross-flow velocity droplet size reaches a plateau^[12,13]^. Our microfluidic device takes advantage of this saturation regime to produce droplets of consistent size while also significantly reducing reagent consumption (requiring only hundreds of μL as compared to tens of mL for traditional systems^[14]^). The device consists of two input channels and an off-the-shelf polycarbonate filter. Each device can be re-used multiple times, takes 35 minutes to fabricate and costs less than $2 (Figure 1b). The geometry of the oil input channel (width; 250 μm, height; 31 μm) results in laminar flow with high shear stress throughout. Each polycarbonate layer, along with the polycarbonate filter, is added sequentially (Supplementary Figure 1) and permanently bonded through thermal fusion for a maximum bond strength of 4.5 MPa (Supplementary Figure 2).

**Figure 1.**
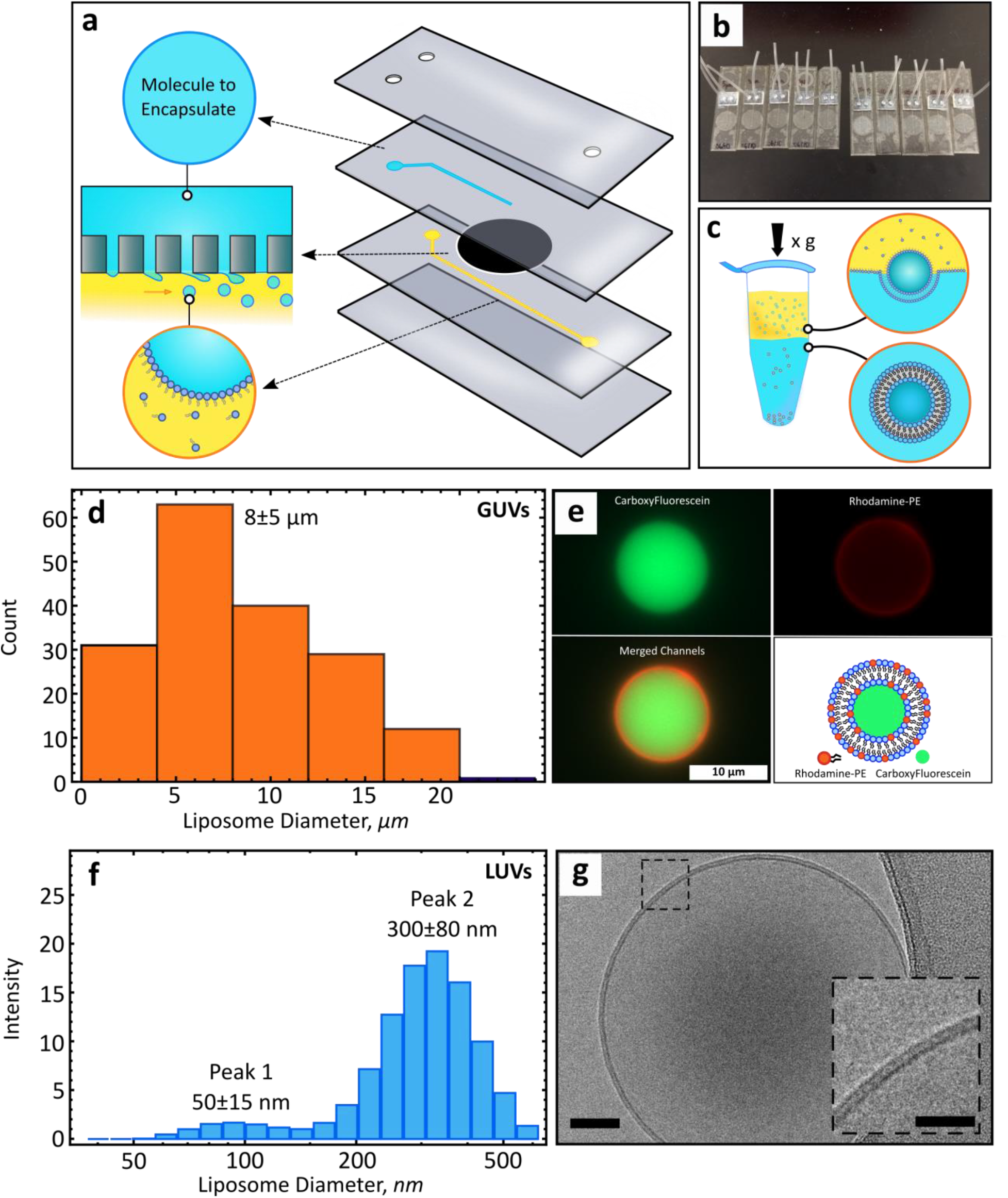
A microfluidic device for generating GUVs or LUVs in two steps. **a**, Schematic of the different layers used to create the final microfluidic device, with an illustration of the droplet formation process used to create micro or nanoscale droplets. Macromolecular targets of interest are pressure driven into the first input channel (blue). Oil solvents saturated with lipids are pressure driven into the second input channel (yellow) forming water-in-oil emulsions. A polycarbonate membrane separates the two channels. **b**, Ten devices can be fabricated in a single session. Each device can be reused multiple times due to low clogging rates. **c**, Phase transfer of lipid-stabilized microscale or nanoscale droplets through a lipid rich interface to form GUVS or LUVS. **d**, Size distribution of GUVs formed with a 5 μm pore membrane (n=176) (mean±s.d.). Measured by optical and fluorescent microscopy. e, Representative image of a GUV encapsulating carboxyfluorescein (CF). Top left, CF alone. Top right, Rhodamine-PE alone (0.5% mol). Bottom left, channel colocalization. Bottom right, schematic of the main features of the GUV. **f**. Size distribution of LUVs formed with a 100 nm pore membrane (n=9) (mean±s.d.). Measured by DLS (Dynamic Light Scattering). **g**, Cryo-EM micrograph of a small unilamellar liposome. Scale bar, 50 nm. Inset scale bar, 25 nm.

The existence of a droplet-size invariant regime was first established theoretically by applying a torque-balance model at the surface of the filter pore^[11]^. Depending on the mode of deformation, we identified droplet-size invariant regions for 5 μm and 100 nm pore polycarbonate filters (Supplementary Figure 3a and 3b). Theoretical modeling, based on the geometric constraints (Supplementary Table 1) used in this work, predicted small droplet diameter variation with a continuous phase flow rate set to 80 μL/min or greater. The existence of this region was confirmed experimentally after the transformation of droplets into liposomes using the PT method (Figure 1c), which has been shown to preserve initial droplet size upon centrifugation^[9,15]^. Dynamic light scattering confirmed droplet-size invariance for a 100 nm filter pore across a variety of flow rates (80 μL/min to 230 μL/min) (Supplementary Figure 3c).

Integration of a 5 μm polycarbonate filter into the microfluidic device lead to the formation of cell-sized lipid vesicles (GUVs) (8±5 μm) (mean±s.d., n=176) (Figure 1d). To visualize these vesicles, carboxyfluorescein (CF) was encapsulated into the lumen of GUVs composed of a simple lipid mixture (POPC/POPS/Cholesterol) with the addition of trace amounts of Rhodamine-labeled PE for lipid bilayer visualization (Figure 1e). The size and distribution of GUVs created here is, on average, smaller and more uniform than GUVs prepared by other microfluidic^[16,17]^ or other methods^[17,18]^ and does not require detergent stabilization^[15]^. Our technique also enables rapid encapsulation of material within the lumen. In addition, by substituting a 5 μm filter for a 100 nm filter we are able to generate organelle-sized lipid vesicles (LUVs) of 300±80 nm (mean±s.d., n=9, Figure 1f). CryoEM enabled direct visualization of LUVs and confirmed the formation of vesicles with intact, unilamellar lipid bilayers (Figure 1g). Compared to spontaneous vesiculation^[19]^ or droplet-extrusion^[19]^, the technique developed here confers improved control over lipid leaflet placement, lamellarity^[20]^, size and distribution.

### Lamellarity, permeability, and solvent contamination

We investigated GUV lamellarity through encapsulation of the dye, carboxyfluorescein. Alpha-hemolysin (aH), a membrane-binding and pore-forming toxin was added to the outside of liposomes. In a series of steps, monomers of aH adsorb on the lipid bilayer, followed by assembly into a heptameric pore that spans the lipid bilayer^[21]^. We observed that aH nanopores successfully assembled within the lipid bilayer and lead to the diffusion of fluorescent molecules out of the vesicle lumen (Figure 2a). Functional alpha-hemolysin insertion demonstrates three things, 1) the lipid bilayer is capable of supporting membrane insertion and pore formation 2) the lipid bilayer is unilamellar and 3) the bilayer remains stable following aH pore formation (Figure 2b).

**Figure 2.**
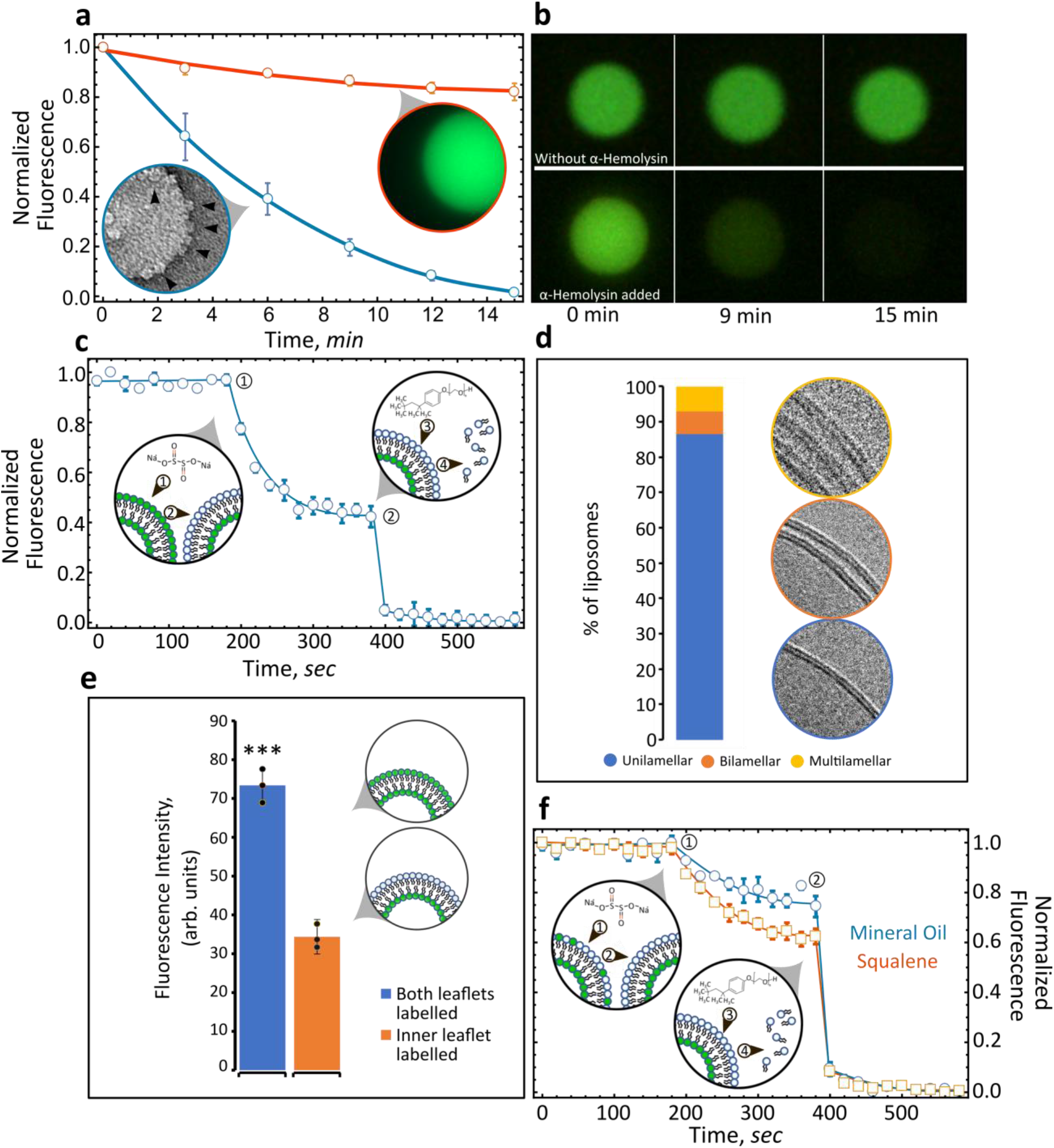
Lipid leaflet lamellarity and content asymmetry of GUVs and LUVs. **a**, GUVs loaded with carboxyfluorescein were monitored over time by fluorescence microscopy. GUVs exposed to alpha-hemolysin (aH) (blue) were compared with control liposomes (red) and the flux of fluorescent molecules from within the lumen monitored over time. Inset: electron micrograph of a liposome studded with aH pores (n=5) (error bars are s.e.m). **b**, Images of GUVs exposed (lower panels) or not exposed (upper panels) to alpha-hemolysin over time (n=5). **c**, Fluorescence intensity and quenching of NBD-PC (both leaflets: POPC/POPS/Cholesterol/NBD-PC) (n=3, error bars are s.e.m). Arrow number indicates, 1) addition of quencher, 2) fluorescence quenching, 3) addition of Triton X-100 and 4) bilayer solubilization and complete quenching. **d**, Liposome lamellarity visualized by cryoEM (lipid composition: POPC/POPS/Cholesterol; n=199). **e**, Fluorescence intensity of liposomes synthesized with NBD-PC present either in both leaflets or a single leaflet (n=3). \*\*\**p* < 0.001. **f**, Fluorescence intensity quenching of NBD-PC deposited exclusively in the inner leaflet. Inner leaflet, POPC/NBD-PC. Outer leaflet, POPS (n=3, error bars are s.e.m). Two different solvents are compared for generating asymmetric liposomes: mineral oil versus squalene

Next, we probed the lamellarity of nanoscale vesicles utilizing two distinctly different assays: fluorescence quenching and cryoEM. First, we utilized sodium hydrosulfite to quench fluorescence from trace NBD-PC lipids. In the vicinity of the fluorophore (NBD) sodium hydrosulfite quenches the fluorescence by reducing the dye. A perfectly symmetric lipid bilayer will result in the reduction of 50% of the signal if the fluorophore (NBD-PC) is evenly distributed between the two leaflets upon exposure to the quenching agent^[9]^. We generated symmetric vesicles composed of POPC/POPS/Cholesterol/NBD-PC (44.8:36.9:17.7:0.5 mol%) and measured NBD-PC fluorescence. For liposomes fabricated with NBD-PC in both the inner and outer leaflets, addition of the quencher reduced total fluorescence by 47% (Figure 2c), suggesting that the dye was evenly distributed in both leaflets and confirming that the majority of liposomes formed by our protocol are unilamellar. To test this notion further, we utilized cryoEM to directly visualize the lamellarity of a population of LUVs. Electron microscopy of vitrified liposomes revealed that approximately 85% of LUVs prepared with this technique are unilamellar with a small fraction of multilamellar (MLV) and multivesicular (MVL) liposomes (Figure 2d, Supplementary Figures 4-5). Significantly, our approach is free of sucrose- or other sugar-containing solutions allowing for cryoEM investigation of lamellarity. Liposomes are fabricated under physiologically-relevant conditions without the need for density-gradients. Sucrose and glucose are routinely utilized for liposome formation and isolation, however high concentrations of sugars are detrimental for cryoEM analysis because they lead to significant reductions in contrast^[22]^.

Next, we generated LUVs with asymmetric lipid leaflets, with an inner leaflet that comprises POPC/NBD-PC and an outer leaflet of POPS alone. We confirmed the total fluorescence of fluorescent lipid headgroups (NBD-PC) present in the two leaflets through fluorescence microscopy. Quantifying the total fluorescence per lipid concentration when only a single leaflet is labeled with NBD compared to both leaflets should reveal approximately a twofold difference. As expected, total fluorescence was 53.2% greater when NBD-labeled PC (0.5 mol%) was included in both stages compared with just the first stage (Figure 2e). A fluorescent quenching assay was performed to quantify the degree of asymmetry with an inner leaflet composed of POPC/NBD-PC and the outer of POPS. Since the lipid bilayer is impermeable to sodium hydrosulfite, a completely asymmetric bilayer should maintain 100% fluorescence upon addition of the quencher. Unexpectedly, we found that oil properties play a role in the degree of asymmetry. Liposomes formed with mineral oil as the solvent, by contrast with squalene, show the greatest level of stable asymmetry, with 79% of the fluorescent intensity protected from the quencher (Figure 2f) (n=3). Imperfect asymmetry is most likely due to the spontaneous movement of lipids from the inner leaflet to the outer leaflet due to either trading of lipids during phase-transfer or flip-flop following fabrication^[23]^. Our findings agree with and reinforce recent publications^[24,25]^, that solvent-type plays a role in the final mechanical and functional properties of lipid membranes.

### Characterizing vesicle utility

Here, we address several outstanding questions regarding the nature of solvent contamination and its impact on the quality and functionality of the lipid membrane. CryoEM imaging enabled us to quantify residual solvent inclusion within lipid bilayers formed by the PT method (Figure 3a). We observed that visible oil lenses within lipid bilayers occur in ~12% of the total liposome population (Figure 3b). When present, solvent lenses average 25 ± 8 nm (mean ± s.d, n = 55) in diameter (Figure 3c). While large oil inclusions are straightforward to characterize (Supplementary Figure 6) we cannot rule out the presence of trace amounts of oil within the lipid bilayer. Interestingly, the observed mineral oil collection in lenses is reminiscent of the fatty acid lenses that form in the endoplasmic reticulum during lipid droplet biogenesis^[26]^. Our methodology can potentially be used to reconstitute and study this phenomenon.

**Figure 3.**
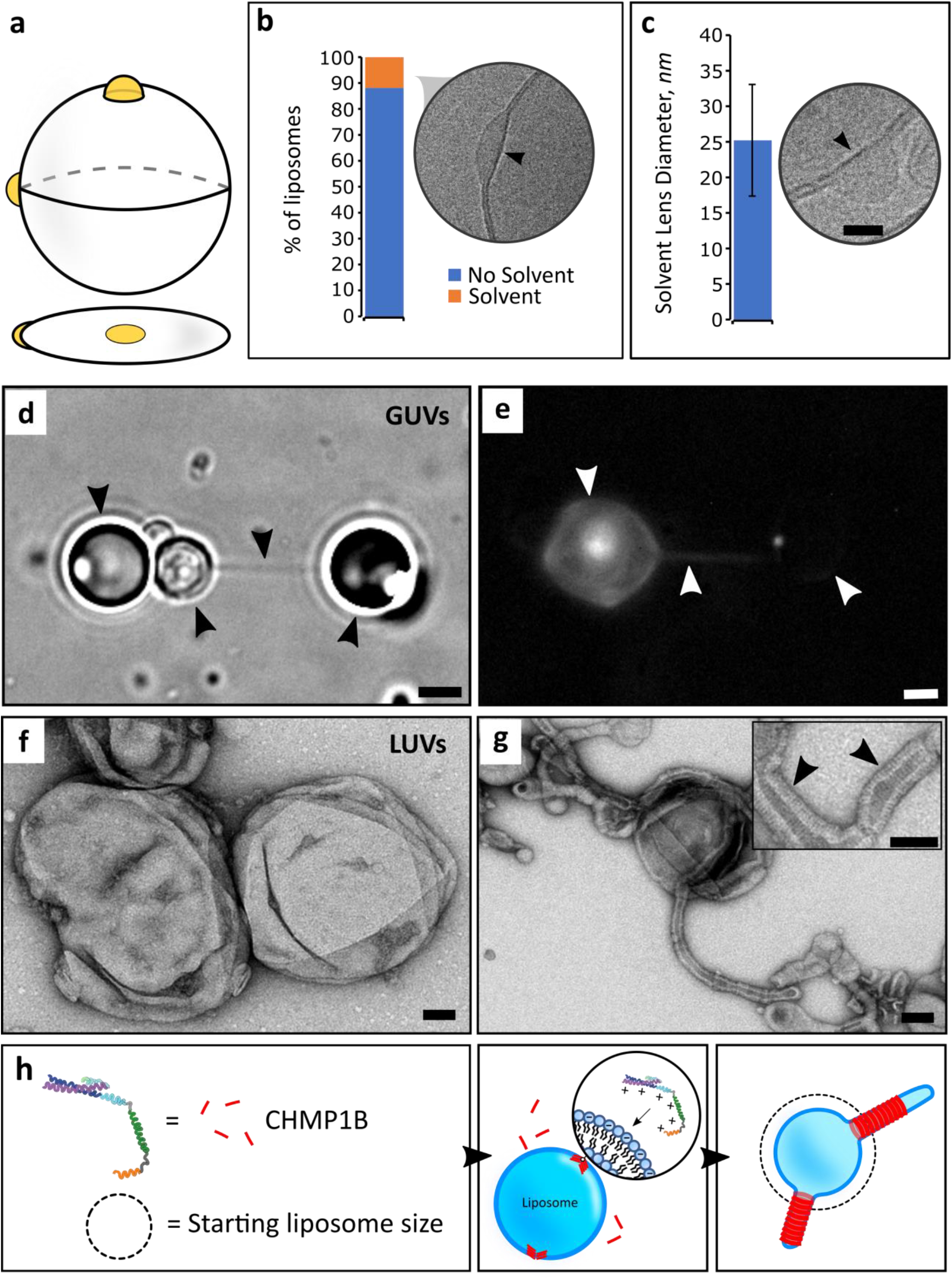
Characterizing oil defects and lipid bilayer properties. **a**, Illustration of oil accumulation as lenses within the lipid bilayer. **b**, Quantification of cryoEM micrographs showing oil lenses within the bilayer (n=1812) (lipid composition, POPC/POPS/Cholesterol). Arrow points to an oil lens located on the outer edge of the liposome. **c**, Oil lens size diameters as observed by cryoEM (n=55) (lipid composition, POPC/POPS/Cholesterol). Arrow indicates typical morphology of solvent lenses. Scale bar, 10 nm. Error bars are s.d. **d**, Optical trap, nanotube pulling experiment. Arrows indicate (from left to right) (5 μm) Silica bead, GUV, nanotube and another Silica bead (POPC/POPS/Cholesterol/Biotin-PE (5 mol%)). Scale bar, 2.5 μm. **e**, Fluorescence micrograph demonstrating incorporation of fluorescent lipids (POPC/POPS/Cholesterol/Biotin-PE/Rh-PE (0.1 mol%)). Arrows indicate (from left to right) a GUV, nanotube and a silica bead. Scale bar, 2.5 μm. **f**, TEM micrograph of liposomes (control), before addition of CHMP1B (lipid composition: POPC/POPS/Cholesterol). Scale bar, 50 nm. **g**, After addition of CHM1B to liposomes. CHMP1B accumulates on the lipid bilayer and subsequently deforms it into lipid tubules. Inset, protein striations (black arrowheads) are clearly visible under negative stain EM. Scale bar, 50 nm. **h**, Cartoon illustration of CHMP1B mediated vesicle remodeling.

Next we utilized nanotube pulling experiments to probe the dynamic response and stability of the lipid membrane. A single lipid tubule pulled from the surface of a GUV requires a constant supply of liquid lipid in order to maintain structural integrity^[27]^. As expected, biotin-coated silica beads conjugated with Traptavidin positioned with an optical trap (Supplemental Figure 7) can be used to pull ~ 50 nm diameter lipid tubules from GUVs incorporating POPC/POPS/Cholesterol with Biotin-PE (Figure 3d) and trace amounts of Rhodamine-PE (Figure 3e). These nanotubes are stable and can extend to tens of micrometers in length.

We also assayed whether vesicles generated using our device are suitable substrates for peripheral membrane-binding proteins. CHMP1B is a human ESCRT-III protein implicated in membrane deformation processes such as recycling tubule biogenesis from endosomes^[6,28,29]^. CHMP1B accumulation on the surface of a lipid membrane requires the presence of negatively-charged lipid headgroups, like POPS. In agreement with previous observations^[6]^, initially spherical liposomes (Figure 3f) are rapidly deformed by electrostatic binding of CHMP1B onto the positive (exterior) surface of liposomes consisting of POPC/POPS/Cholesterol (Figure 3g). CHMP1B quickly saturates the liposomal surface, inducing strong positive curvature along the tubule axis. The outer protein shell is clearly visible under TEM (Figure 3g) demonstrating that CHMP1B is able to efficiently bind, stabilize and deform these vesicles (Figure 3h). This overlap of the behavior we observed for CHMP1B with previously published results reinforces the idea that the GUV’s formed by our microfluidic device are biologically relevant.

### Encapsulation of synthetic and organic macromolecules

Liposomes have been used extensively as cargo carriers because the inner lumen of the liposome is protected from the exterior environment by the lipid bilayer. By including different cargoes in the first stage of emulsion generation, we encapsulated two cargoes with different properties. An advantage of our method is the ability to create GUVs and LUVs incorporating diverse solution conditions, e.g. high ionic strength buffers (> 350 mM NaCl), for encapsulation of target proteins that would otherwise polymerize or assemble under physiological conditions.

For encapsulation we first loaded LUVs with computationally designed, self-assembling 25 nm dodecahedral nanocages (Figure 4a)^[30]^. These recently reported^[30]^ nanocages hold great promise for drug delivery and synthetic biology applications, including as cargo carriers^[31]^, as imaging probes^[32]^ and as scaffolds for vaccine design^[33]^. Initially, we purified and visualized 25nm diameter aldolase nanocages^[30,34]^ by negative stain TEM (Supplementary Figure 8). We then demonstrated nanocage encapsulation within vesicles manufactured in our microfluidic device by 3 methods: 1) We observe that aldolase activity is sequestered within liposomes, but can be subsequently released upon the addition of a detergent (Figure 4b); 2) TEM imaging revealed controlled nanocage release from detergent-exposed liposomes (Supplementary Figure 9 and Supplementary Figure 10a); 3) Finally, direct visualization by cryoEM demonstrated that the protein nanocages inside the synthesized liposomes matched the expected 25 nm size and assembled architecture of the designed structure (Figure 4c,d and Supplementary Figure 10b)^[30]^. In addition to these dodecahedral nanocage assemblies, we also encapsulated GFP (Green Fluorescent Protein) within both GUVs and LUVs, as determined by fluorescence microscopy (Figure 4e-f). Developing efficient means by which biological macromolecules can be encapsulated and protected while maintaining their structure and function will prove vital for future applications.

**Figure 4.**
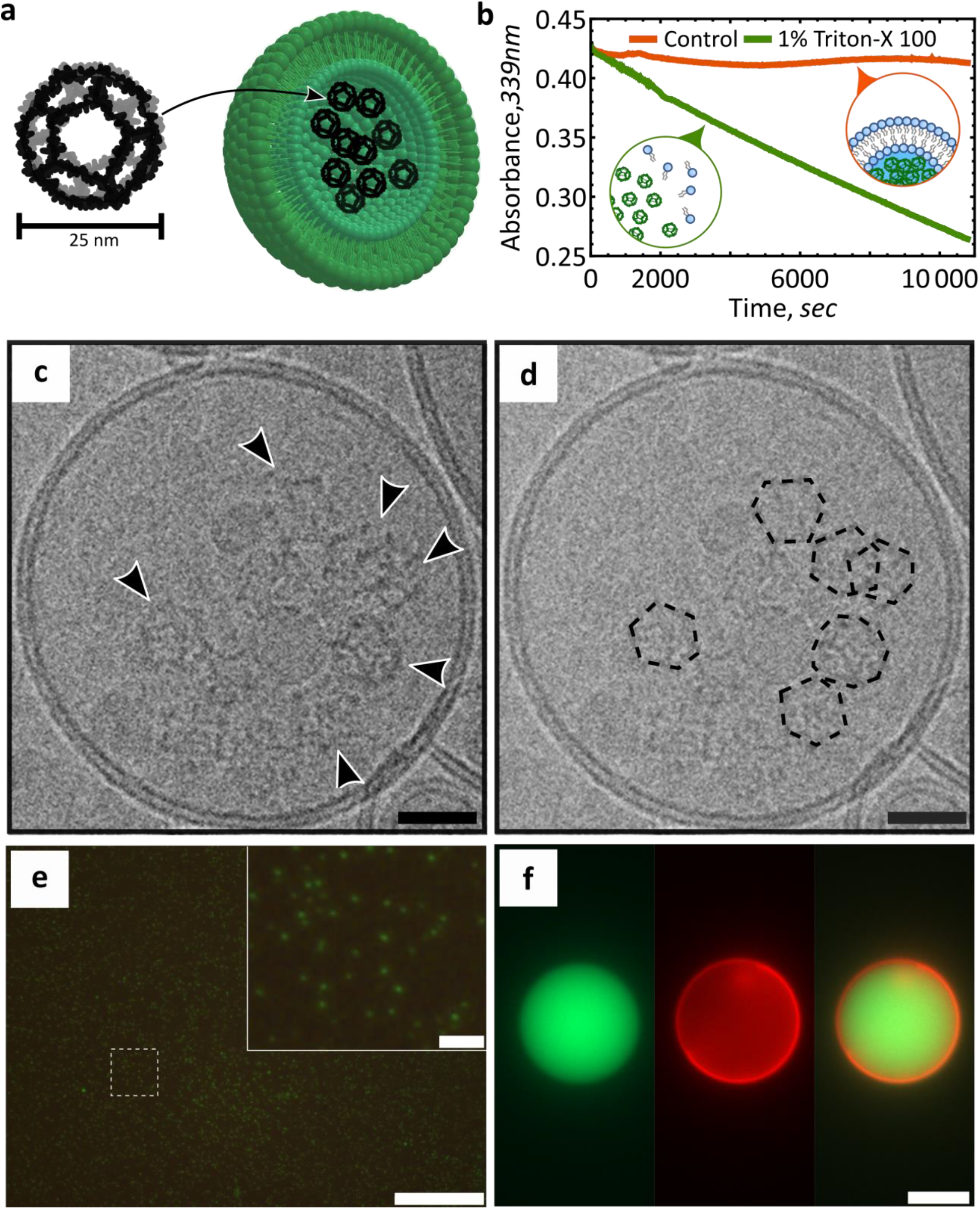
Cargo encapsulation within LUVs or GUVs. **a**, Cartoon representation of the designed dodecahedral nanocage structure and encapsulation within a liposome. Lipid bilayer not to scale. Adapted from^[30]^. **b**, Protection and release assay of designer nanocages. Aldolase nanocages are protected within liposomes (orange) and are exposed following addition of 1% Triton X-100 (green) (lipid composition: POPC/POPS/Cholesterol; n=3). **c** and **d**, CryoEM micrograph of a nanocage loaded liposome (lipid composition: POPC/POPS/Cholesterol). Arrows indicate nanocages. Scale bar, 25 nm. Nanocages are outlined to aid with visualization. **e**, Fluorescence micrograph of GFP encapsulated within LUVs (lipid composition: POPC/POPS/Cholesterol). Scale bar 10 μm; inset scale bar, 1 μm. **f**, Fluorescence micrograph of GFP encapsulated within GUVs (lipid composition: POPC/POPS/Cholesterol/Rhodamine-PE). Scale bar, 10 μm.

## Conclusion

We have developed a simple, inexpensive and modular microfluidic approach for the formation of size-controlled, lipid-content controlled, lumen-content controlled, and asymmetric liposomes that comprise, on average, a single phospholipid bilayer. Utilizing cross flow emulsification theory, modified for the geometric properties of microfluidic channels, we identified and experimentally verified the existence of droplet-size flow invariant regions. When operating within this regime, use of the appropriate polycarbonate filter leads to the creation of either nanoscale or microscale vesicles. CryoEM studies, enabled by sugar-free solutions, revealed the extent and size of oil inclusions within liposomes. We show that our vesicles are able to support a variety of complex phenomena such as nanotubule formation and lipid bilayer remodeling by an ESCRT-III protein. The lipid vesicles created here proved remarkably stable to a host of different chemical assays, demonstrating effective encapsulation and protection of different cargoes.

## Experimental Section

### Materials

Polycarbonate, clear, laser markable film (SD8B94) at a thickness of 50 μm was purchased from SABIC. PCTE, hydrophobic, 0.1 and 5 μm polycarbonate membranes were purchased from Sterlitech Corporation (PCTF0113100, PCTF5013100). Tygon 0.02×0.06" tubing was purchased from Cole-Parmer. UV curable adhesive (3106) was purchased from Loctite. Nanosep 10 kDa MF Centrifugal Devices (Spin columns) were obtained from VWR. Wellplates (384 well, black bottom, polystyrene) were purchased from Corning. Axygen 2 mL microtubes (MCT-200-L-C) were purchased from Axygen, Inc. Continuous carbon film grids (Formvar/Carbon Film (FCF-200-Cu)), Quantifoil holey carbon grids (2 μm hole size, 2 μm spacing, 200 mesh) and ultrathin carbon supported by a lacey carbon film on a 400 mesh copper grid (# 01824) were obtained from Ted Pella Inc. Slide-A Lyzer MINI Dialysis Devices (10K, 88401) were purchased from Thermo Scientific.

### Nanoemulsion and LUV formation

Lipid handling and preparation procedures followed those of previously published protocols^[35,36]^. Phospholipids (POPC/POPS/Cholesterol (45.3:36.9:17.7 mol%)) (unless otherwise stated, this lipid ratio was used for all formulations and is the standard lipid mixture) stored in chloroform were dispersed in mineral oil at a final lipid concentration of 5 mM. Glass vials were placed into an oven overnight to evaporate any chloroform. The oil-lipid mixture was further diluted to 2 mM using mineral oil. A 1 ml syringe loaded with 1xPBS was loaded into a syringe pump (KD Scientific 200) and driven at a constant volumetric flow rate of 4 μl/min through the upper channel. Another syringe pump (KD Scientific 220) was loaded with a 3 ml syringe. 130 μl of emulsion was removed from the outlet of the device and added to a 2 ml microtube. Emulsions were placed into a fridge at 4°C for 20 minutes to allow the lipids to equilibrate at the droplet interface. Concurrently, 130 μl of 1xPBS was added to the bottom of a 2 ml microtube. Subsequently, 170 μl of the oil-lipid mixture was placed on top of the buffer and allowed to equilibrate for 20 minutes at room temperature. Finally, emulsions were distributed between vials and centrifuged (4°C for 10 minutes, 20,000 x g).

### GUV formation and carboxyfluorescein encapsulation

Following the same protocol as outlined for LUVs, microemulsions were synthesized by keeping the dispersed phase constant at 5 μL/min and the oil flow rate constant at 120 μL/min. Following incubation at room temperature, emulsions were added to the capture vial and centrifuged (10 minutes at 9,000 x g). Carboxyfluorescein (350 mM NaCl, 50 mM Tris-HCl pH 7.0, 5% (w/v) Glycerol and 5 mM β-mercaptoethanol) was encapsulated within GUVs at a final concentration of 50 μm.

### Liposome leaflet asymmetry assay

The inner lipid leaflet was composed of POPC/NBD-PC (99.5:0.5 mol.%). The outer leaflet was composed exclusively of POPS (100 mol.%). Emulsion preparation and collection followed previously described steps except for the addition of NBD-PC to the emulsion formation phase. Unlabeled POPS oil-lipid mixture was diluted with mineral oil (1:1). 170 μl of this mixture was placed on top of 130 μl of 1xPBS buffer in a 2 ml microtube and allowed to equilibrate for 20 minutes at room temperature. Emulsions were centrifuged at 300 *g* for 10 minutes at room temperature. Following previously published protocols^[37]^, 10 μl of unconcentrated liposome solution and 10 μl of buffer were added to one well and 20 μl of buffer was added to another. After setting the baseline, 0.5 μl of 1 M sodium hydrosulfite prepared in 1xPBS was added to the sample followed by the addition of 2 μl of 10% Triton X-100.

### Aldolase enzyme activity assay

The 2-keto-3-deoxy-6-phosphogluconate (KDPG) aldolase activity of the I3-01 nanocage domain was monitored using an L-lactic acid dehydrogenase (LDH)-coupled assay^[34,38]^. 10 μL of nanocage-encapsulated liposomes were mixed with 90 μL of 1xPBS, 0.1 mM NADH, 0.11 U/μl 1 LDH, 1 mM KDPG and either including or omitting 1% Triton X-100. Loss of absorbance at 339 nm owing to oxidation of NADH was monitored using a Synergy Neo2 microplate reader. At least three replicates of the aldolase activity assay were performed.

### CryoEM imaging

Liposomes loaded with 1xPBS were prepared as described above and 3 ml of the sample was concentrated to 100 uL. For nanocage-encapsulated liposomes, 200 μg/ml nanocages were loaded into liposomes and concentrated from 1 ml to 100 μL. For electron cryo-microscopy, 3.5 μl of these concentrated samples were applied to either glow-discharged Quantifoil holey carbon grids (2 μm hole size, 2-4 μm spacing, 200 mesh) or Ultrathin carbon film on holey carbon grids (400 mesh), blotted (6.5-8 seconds, −1 mm offset) and plunge-frozen in liquid ethane using a Vitrobot Mark I (FEI). Electron cryo-micrographs were collected following low-dose procedures at liquid nitrogen temperature on a Tecnai TF20 operating at 200kV using a Gatan 626 side-entry cryo-holder. Movies were recorded using a K2 Summit direct detector (Gatan, Pleasanton, CA) in either counting or super-resolution mode at a corrected magnification of 41,911x, corresponding to a physical pixel size of 1.193 Å, and at dose rates of ~7 e-/pixel/sec at the specimen. SerialEM^[39]^ was used to facilitate low-dose imaging and semi-automated data collection, and each movie was recorded as a stack of 40 subframes, each of which was accumulated for 0.2 s, totaling ~39 e-/Å^2^ at the specimen. Frames were aligned and summed by using MotionCor2^[40]^.

## Acknowledgments

We thank David Belnap at the University of Utah Electron Microscopy Core Laboratory for help with cryoEM image acquisition and processing, Joerg Votteler for guidance on aldolase assays as well as Marc Porter and Hamid Ghandehari for access to the Malvern Zetasizer. We thank Wes Sundquist for numerous discussions and critical reading of the manuscript. AF acknowledges funding by a Faculty Scholar grant from the Howard Hughes Medical Institute, the Searle Scholars Program, NIH grant 1DP2GM110772-01, the American Asthma Foundation and the Chan Zuckerberg Biohub. JM and AF acknowledge funding by the National Institutes of Health (P50 GM082545 and R01 GM127673-01). MV acknowledges funding by National Science Foundation (ENG/CMMI #1563280).

## Author contributions

V.R., M.V., J.M. and A.F. designed the experiments. V.R., J.M., and A.S. performed the experiments. V.R. analyzed the data. B.K.G. contributed to the device design, fabrication and characterization. V.R., J.M. and A.F. wrote the paper. All authors had direct input on experimental results and the preparation of the manuscript.

## Additional information

Supplementary information is made available.

