## Supporting Information for "A tunable microfluidic device enables cargo encapsulation by cell-or organelle-sized lipid vesicles comprising asymmetric lipid monolayers"

**Corresponding authors (email)**

**Methods**

**Reagents**. POPC (1-palmitoyl-2-oleoyl-sn-glycero-3-phosphocholine), POPS (1-palmitoyl-2-oleoyl-sn-glycero-3-phospho-L-serine (sodium salt)) and Cholesterol were purchased from Avanti Polar Lipids, Inc. Triton X-100, Sodium Hydrosulfite ($\mathrm{Na_2S_2O_4}$), α-Hemolysin from *Staphylococcus aureus*, KDPG (2-keto-3-deoxy-6-phosphogluconate) and NADH (β-Nicotinamide adenine dinucleotide) were purchased from Sigma-Aldrich, Inc. LDH (L-lactic acid dehydrogenase) was purchased from Roche. Purified Green Fluorescent Protein was a generous gift from Matt Lalonde.

**Microfluidic device fabrication.** The microfluidic device architecture was designed in AutoCAD 2014 and uploaded to a Versa Laser, VLS 3.60 (Universal Laser Systems). Laser power was set to 1.90% with a speed of 7.50%. Once cut, each half of the device was aligned and thermally bonded using a heat press (JetPress 12, Geo Knight and Co., Inc.) at 150°C for 2 minutes. Channel width was 200 µm and height was determined by the thickness of the polycarbonate sheet (50 µm). Using a light camera (AM7115MZ 5MP,Dino-Lite), channel width was measured at 250 µm with a height of 31 µm after thermal bonding. Subsequently, a membrane was added to one of the bonded halves and restrained using spot welding (light touch with the tip of a soldering iron). The two halves of the microfluidic device were then thermally fused (150 C for 2 minutes). Acrylic ports were adhered to the top of the device to allow for placement of tubing for fluid delivery. Tygon inlet tubes were affixed to the acrylic ports via UV curable adhesive (long wavelength, 30 minutes curing time).

**Size-invariant flow model**. Droplet detachment is driven by a balance of forces at the outlet of the membrane pore[1]. This is only valid under the following assumptions, laminar fluid flow, particle either perfectly spherical or highly deformed upon detachment, straight membrane pores and no interfacial tension gradients. The balance of forces is written as follows,

$$F_{RETAINING} = F_{DETACHMENT} \tag{1}$$

The interfacial tension force ($F_{\mathrm{IT}}$) is the major retaining force and is formed due to the dispersed phase (solution) adhesion around the opening of the membrane pore. Interfacial tension forms over an area between the dispersed and continuous phase,

$$F_{\mathrm{IT}} = \pi D_P \gamma \tag{2}$$

where $D_P$ is the pore diameter and $\gamma$ is the interfacial tension between the dispersed phase and the continuous phase.

Droplet detachment is driven by several forces. The most significant force is the drag force ($F_{\mathrm{DF}}$) which is formed by flowing oil over the surface of the membrane. The higher the shear stress, the greater the ability to control droplet size.

$$F_{\mathrm{DF}} = 3\pi f_c k_c \rho_{oil} u^2 {D_D}^2 \qquad (3)$$

where $f_c$ is the Fanning correction factor, $k_c$ is the wall correction factor taken as $k_c = 2.6$[2], $\rho_{oil}$ is the density of the oil-lipid mixture, $u$ is the average channel velocity and $D_D$ is the droplet diameter.

The static pressure force is the second significant force ($F_{\mathrm{ST}}$). It is the pressure difference between the dispersed phase and continuous phase at the membrane surface. It is also known as the Young-Laplace force. Once this force is overcome the droplet starts to contract at the neck of the pore.

$$F_{\mathrm{ST}} = \frac{\gamma}{D_D} \pi {D_P}^2 \qquad (4)$$

The dynamic lift ($F_{\mathrm{DL}}$) and buoyancy ($F_{\mathrm{BG}}$) forces are typically more significant in systems where large droplets are generated. Dynamic lift results from the parabolic velocity profile (asymmetric velocity distribution) within the channel while the buoyancy force is driven by the density difference between the continuous phase and the dispersed phase,

$$F_{\mathrm{DL}} = 2\frac{\tau^{1.5}\rho_{\mathrm{oil}}^{0.5}}{\mu_{\mathrm{oil}}} {D_D}^3 \qquad (5)$$

$$F_{\mathrm{BG}} = \frac{1}{6}\pi g(\rho_{\mathrm{oil}} - \rho_{\mathrm{solution}}) {D_D}^3 \qquad (6)$$

where $\tau$ is the wall shear stress, $\mu_{\mathrm{oil}}$ is the viscosity of oil, $\rho_{\mathrm{solution}}$ is the density of the dispersed phase and $g$ is gravity. With,

$$\tau = \frac{u^2 \rho f_c}{2} \qquad (7)$$

$$f_c = \frac{24(1 - 1.3553a_c + 1.9467a_c^2 - 1.7012a_c^3 + 0.9564a_c^3 - 0.2537a_c^4)}{Re} \qquad (8)$$

where $f_c$ is the fanning friction factor for rectangular channels[3], $a_c$ is the channel aspect ratio and $Re$ is the Reynolds number with the characteristic length scale defined as the hydraulic diameter using the average channel velocity.

By taking a torque balance around the neck of the droplet at the membrane pore outlet we arrive at an algebraic expression that describes the balance of forces for a spherical droplet moving away from the surface of the membrane,

$$[F_{IT} + F_{DL} + F_{BG} - F_{ST}]\frac{D_P}{2} = F_{DF}\frac{D_D}{2} \tag{9}$$

If the droplet is deformed along the length of the surface of the membrane, the droplet size can be approximated via the following modification to the above equation,

$$[F_{IT} + F_{DL} + F_{BG} - F_{ST}] = F_{DF} \tag{10}$$

The direction of forces needs to be considered based on how the dispersed phase is driven through the membrane. This force balance results in a polynomial expression for $D_D$, which can be solved numerically.

**Liposome size and distribution.** GUVs were visualized with fluorescent microscopy (Zeiss Axion-1) and measured using ImageJ. LUVs were synthesized at a constant dispersed flow rate of 3 µL/min and a constant continuous phase of 150 µL/min. LUV size was measuring using Dynamic Light Scattering (DLS) (Malvern Nano-ZS). One hundred microliters of LUVs were added to the bottom of a Malvern disposable capillary cell (DTS1070). Attenuation and number of runs were set to automatic. Every 100 µL of liposomes were measured 3 times with at least 12 runs per measurement.

**Pressure measuring.** A 1 mL syringe, loaded with de-ionized water was placed into a 2-channel syringe pump. A 3 mL syringe loaded with mineral oil was placed into a 10-channel syringe pump. High pressure tubing was attached to the 3 mL syringe with the other end attached to a 1600 PSI pressure transducer (MK-1600). The output line from the transducer was attached to the device. Pressure drop across the device was monitored via an interface written in LabVIEW. Maximum pressure was measured by blocking the oil inlet and outlet of two different versions of the device, one with a membrane and another without. De-ionized water was driven into the device at 100 µL/min via a high-pressure pump (LC-10AD, Shimadzu) until failure.

**Liposome leaflet symmetry assay.** Liposomes were fabricated in the same manner as described in the liposome leaflet asymmetry assay except for the addition of NBD-PC (0.5 mol.%) to the standard lipid composition (POPC : POPS : Cholesterol: NBD-PC (44.8 : 36.9 : 17.7: 0.5 mol%)). Emulsions were created at an oil flow rate of 150 µL/min. After centrifugation, 1 mL of the liposome solution was further concentrated using a spin column. The final working volume was 50 µL. Fluorescence quantification was carried out using a microplate reader (Synergy Neo2 Multi-Mode reader (Biotek)). Excitation was set to 460 nm with emission at 534 nm with temperature averaging 22.2 °C. 5 µL of liposomes and 5 µL of 1xPBS buffer were added to one well of a 384 well plate and another 10 µL of 1xPBS added to another well. After setting the baseline, 2 µL of 1 M sodium hydrosulfite (quencher) freshly prepared in 1xPBS was added to each well. The quencher was prepared fresh on the day of use. After the signal had stabilized, a further 2 µL of 10% Triton X-100 prepared in 1xPBS was added to each well.

**Liposome remodeling by CHMP1B.** CHMP1B protein was purified[4] and added to liposomes[5] as previously described. Briefly, liposomes encapsulating 150 mM Sodium Chloride (NaCl) were created using our microfluidic approach (above). CHMP1B was directly added to 25 µL of liposomes (10 µM final protein concentration) in a total volume of 100 µl and incubated in a medium strength ionic buffer (50 mM Tris pH 8.5, 100 mM NaCl) overnight at room temperature. Assemblies were concentrated 10-fold by low speed centrifugation (2,300 x *g*, 5 min) and re-suspended in the medium strength ionic buffer for negative staining.

**Negative stain electron microscopy.** Liposomes were prepared for negative stain transmission electron microscopy (TEM) following established protocols[6]. Specifically, 4 µl of sample was applied to glow-discharged (PELCO EasiGlow, 15 mA, 0.39 mBar, 25 seconds) continuous carbon film grids and stained with saturated uranyl acetate. Negatively stained grids were imaged on a JEM-1400 (JEOL) microscope equipped with a LaB6 filament and operated at 120 kV. Images were recorded at a nominal magnification of 10,000-30,000x using a charge-coupled device camera (Gatan Orius SC1000B).

**Nanocage purification and encapsulation.** Purified aldolase nanocages (EPN-01; Myr-I3-01-p6) were synthesized as described previously[7] with additional purification steps. Specifically; 1) 0.05% equivalent (v:v) of 10% polyethyleneamine (PEI) was added after cell lysis and nucleic acid precipitates removed by centrifugation and 2) the protein was flown over a Q column to remove contaminants before final polishing by gel filtration chromatography in 25 mM Tris pH 8.0, 150 mM NaCl, 5 mM EDTA. EPN-01 nanocages at 100 µg/ml were loaded into liposomes (following the asymmetry protocol except with the same inner and outer leaflets; inner and outer leaflets: POPC/POPS/Cholesterol (44.8 : 36.9 : 17.7 mol%). Mineral oil was combined with olive oil (1:1) in order to form the outer leaflet.) and the resulting nanocage-encapsulated liposomes concentrated from 1 ml down to 100 µL.

**Detergent-controlled nanocage release**. Nanocage-encapsulated liposomes were incubated at room temperature for 20 minutes in the absence or presence of 15 mM Octyl-glucopyranoside (A.G. Scientific, Inc.). Released nanocages were visualized by negative stain TEM (see above) and quantified as follows: Each grid square (n=4), was screened for 40 minutes without retracing. A statistical analysis using ANOVA (Analysis of Variance) and a Tukey HSD (honest significant difference) test was used to quantify differences between samples.

**Alpha-hemolysin Incorporation.** Liposomes were loaded with a high ionic strength buffer (350mM NaCl, 50 mM Tris-HCl pH 7.0, 5% (w/v) Glycerol and 5 mM β-mercaptoethanol) and prepared as described. Alpha-hemolysin was reconstituted in the same buffer and added to the outside of liposomes (final concentration 10 µg/mL). The combined solution was transferred to a 500 µL dialysis vial placed on top of a 1 L de-ionized water-filled beaker. After two and a half hours, the solution was spun down (5000 x g, 5 minutes) and sediment collected for negative staining. For visualizing fluorescein leakage, we added 5 µL of GUV solution onto a glass slide. Next, we added 1 µL of 500 µg/mL solution of alpha-Hemolysin. A cover slip was placed immediately over the droplet and visualized.

**GFP Encapsulation in LUVs.** His-tagged GFP in high ionic strength buffer (350mM NaCl, 50mM Tris-HCl pH 7.0, 5% (w/v) Glycerol, 5mM β-mercaptoethanol) was mixed with 200 mM Sucrose. 100 mM Sucrose and 100 mm Glucose were added to the capture vials. Lipids in oil were prepared as described (inner and outer leaflets: POPC/POPS/Cholesterol (44.8 : 36.9 : 17.7 mol%)). Emulsions were equilibrated at room temperature for 20 minutes. Emulsions were then added to already stabilized vials and spun at room temperature (16,000 x g for 15 minutes). An Epi-Fluorescence microscope (BX50WI, Olympus) with a FITC filter (excitation 490 nm, emission 520 nm) using an 50x oil-immersion lens (LMPlanFl, 50x/0.50, Olympus) was used for GFP loaded liposome visualization. 10 µl of liposomes were placed onto a glass slide (VWR) with a cover glass (18x18-1.5 Fisherbrand, Fisher Scientific) placed on top. A drop of oil (Immersion oil type-f, Olympus) was then added to the cover glass.

**GFP Encapsulation in GUVs.** GUVs loaded with His-GFP utilized the same buffer minus the Sucrose and Glucose. Rhodamine-PE (0.1 mol%) was added to the lipid mixture to help visualize the lipid bilayer. After equilibrating, droplets were added to the oil-water capture buffer vial and spun down at 9,000 x g for 10 minutes. GUVs were then collected and imaged on a fluorescent microscope at 40x oil-immersion lens.

### Supplementary figures

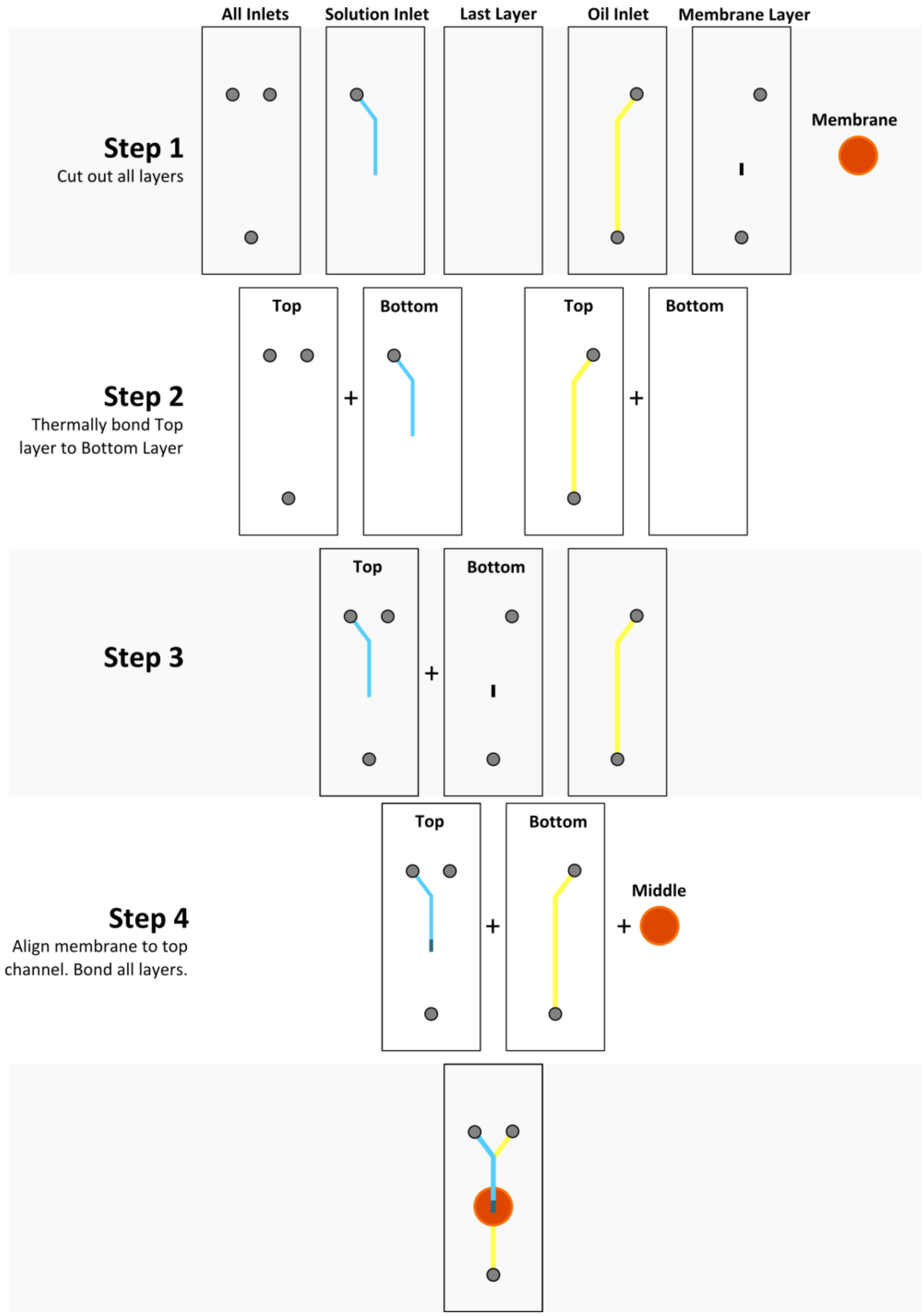

**Figure 1 | Fabrication work flow**
Step by step description of channel bonding sequence and membrane integration.

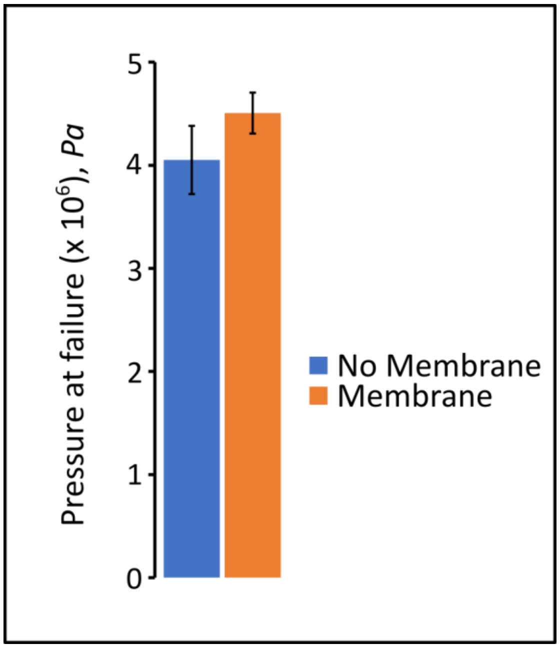

**Figure 2 | Maximum pressure before device failure.**

Two configurations were created, one device without a membrane (left) and another with (right). Common modes of failure included leaking at the inlets and polycarbonate rupture (formation of a small hole).

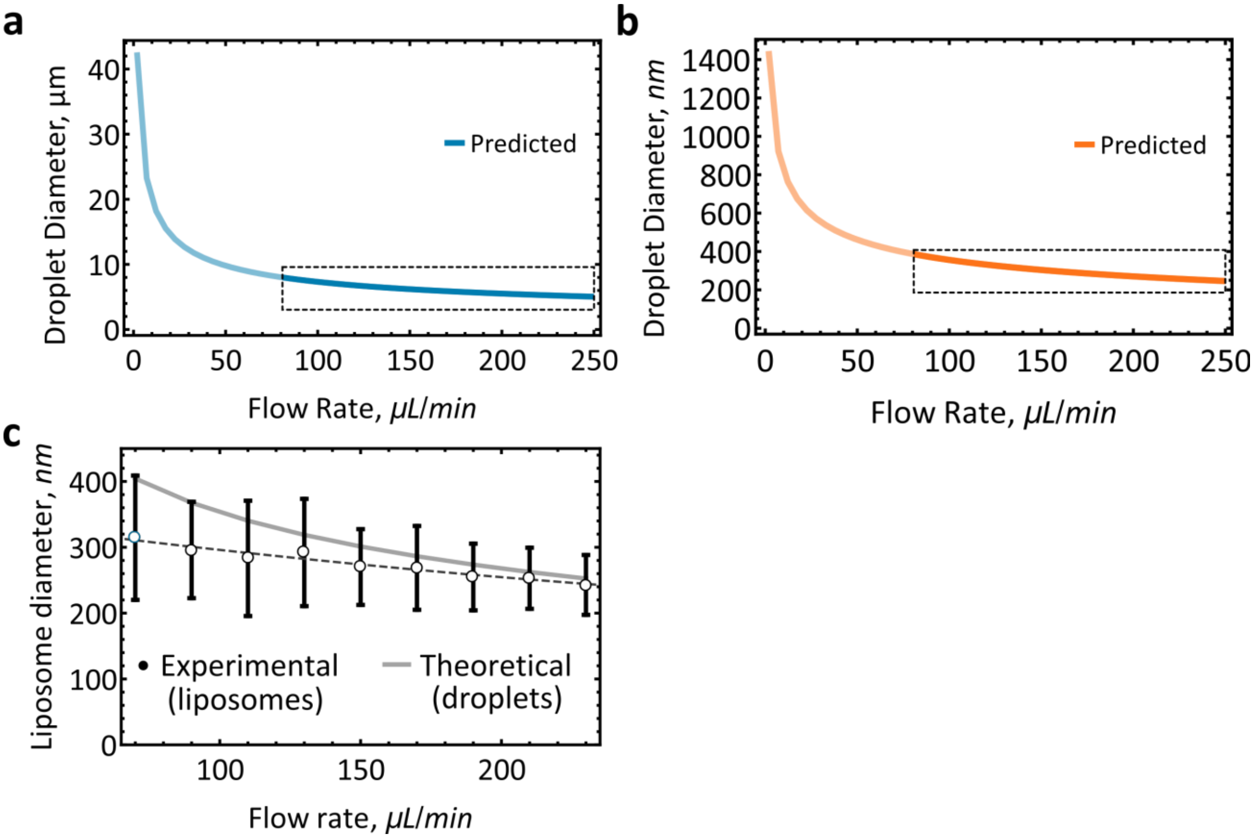

**Figure 3 | Size-invariant flow region.**

**a**, Droplet size prediction based on a 5 µm pore membrane. **b**, Droplet size prediction based on a 100 nm pore membrane. Dotted box highlights the area of flow rate independent droplet formation. **c**, Experimental results of liposome diameters when using a 100 nm pore membrane. Flow rate based on the invariant-size prediction from part b. Mean±s.d.

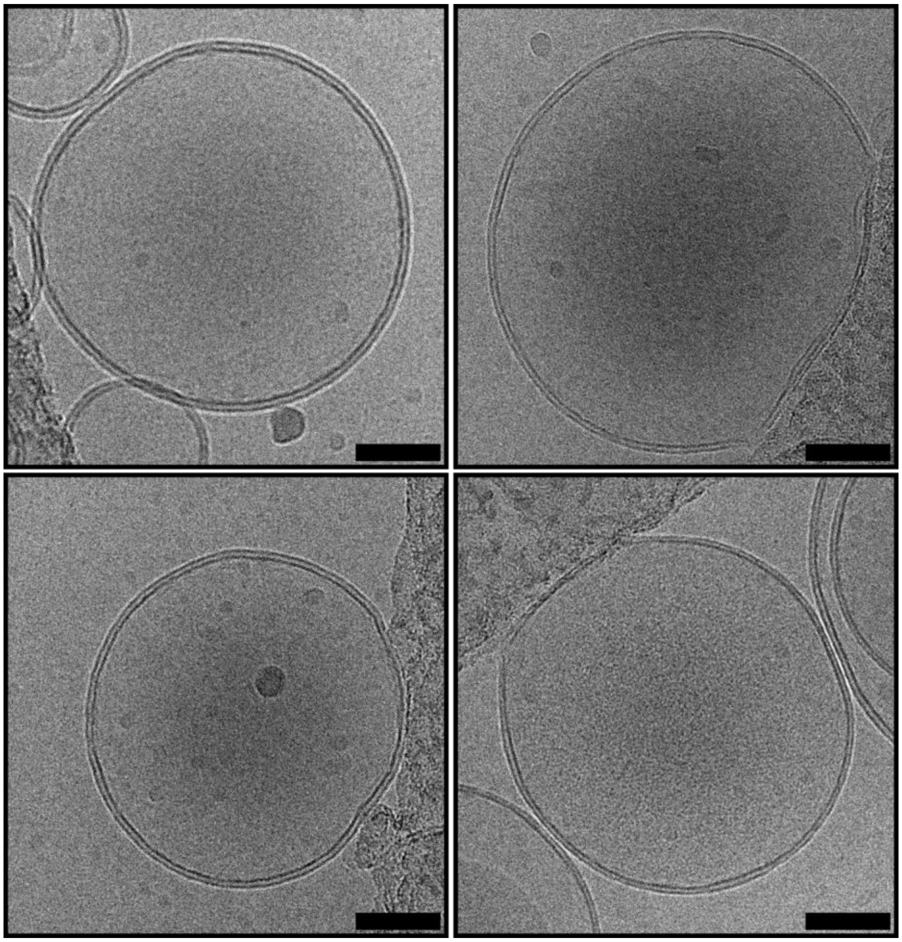

**Figure 4 | Unilamellar Liposomes.**

Representative gallery of cryoEM micrographs of unilamellar liposomes. Scale bar, 50 nm.

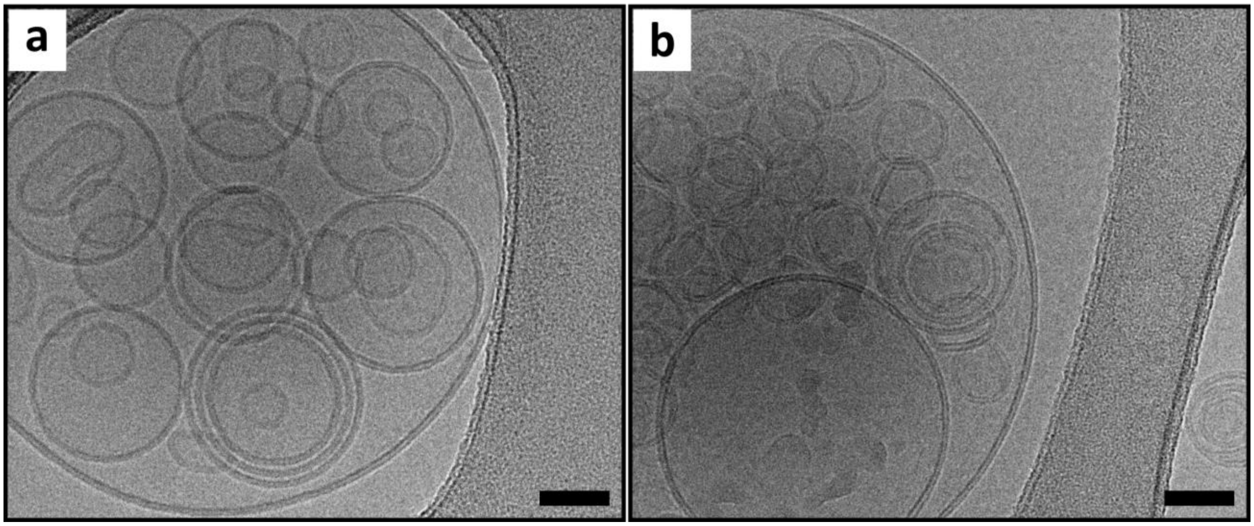

**Figure 5 | Multivesicular Liposomes.**

Representative gallery of CryoEM micrographs of multivesicular liposomes. Scale bar, 50 nm.

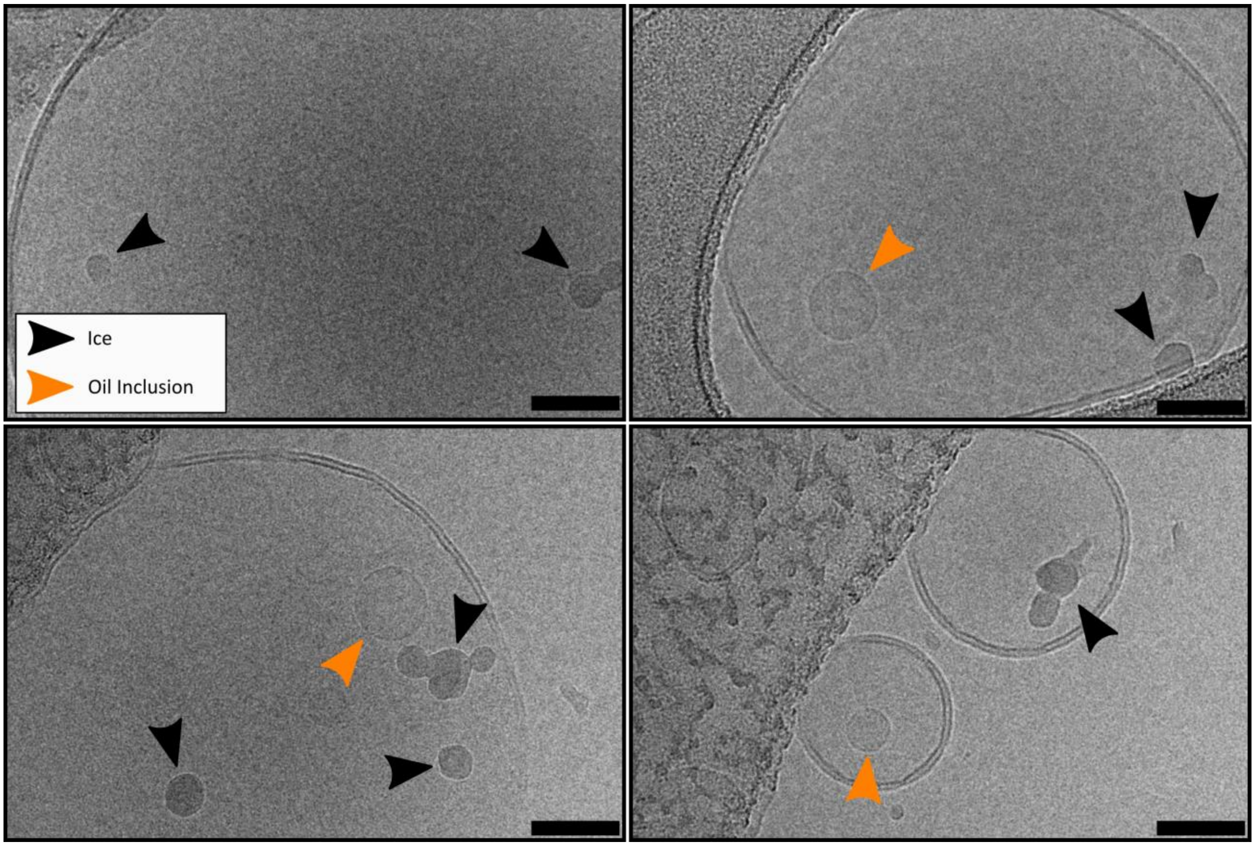

**Figure 6 | Discriminating oil lenses from other contaminants.**

Representative gallery of apparent oil lenses versus other common ice contaminants with regards to irregular boundaries and contrast. Black arrowheads indicate crystalline ice and other cryoEM contaminants, orange arrowheads indicate apparent oil lenses. Scale bars, 50 nm.

a

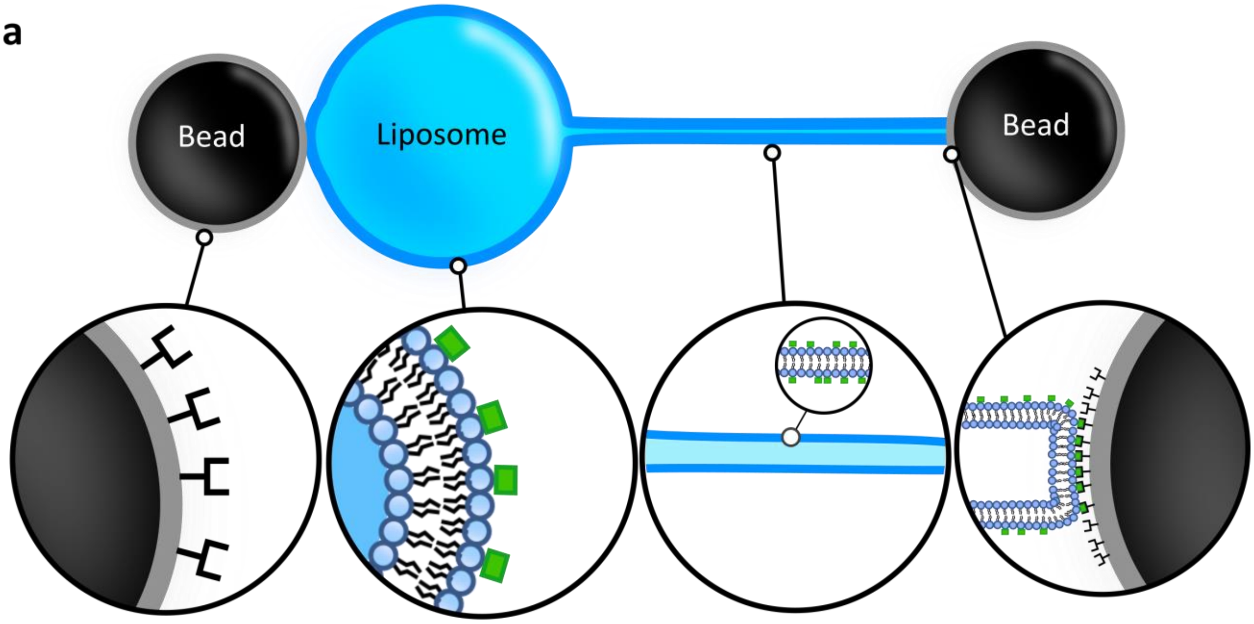

**Figure 7 | Optical-trap nanotube pulling.**

**a**, Schematic of the setup. Silica microbeads are conjugated with Biotin-Traptavidin. Liposomes incorporate POPC/POPS/Cholesterol/Biotin-PE (5 mol%). Biotin-PE is shown as green squares. One bead is used to immobilize the GUV while the other bead is used to pull a nanotube.

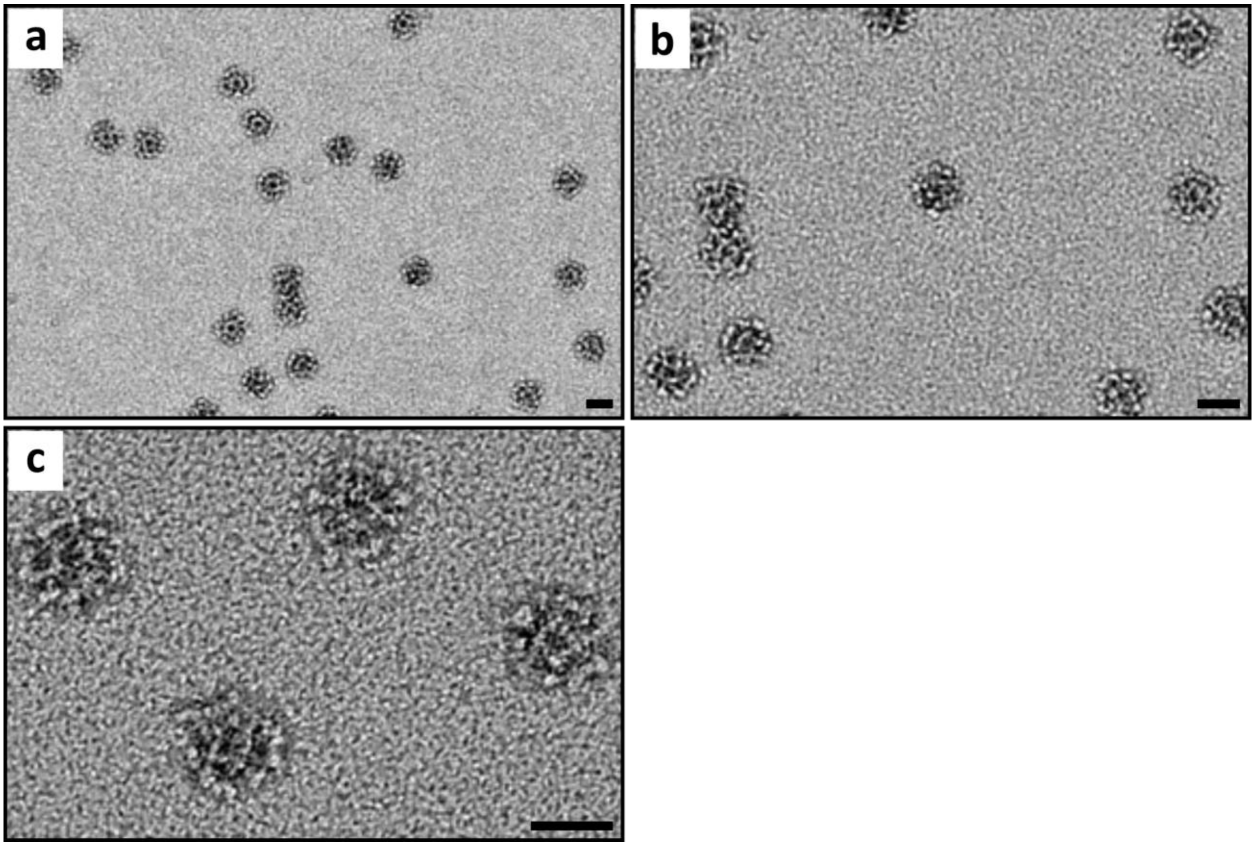

**Figure 8 | Purified nanocages**

Transmission electron micrographs of purified nanocages **a**, low magnification view. **b**, medium magnification view of the same area. **c**, high magnification view. Scale bars, 25 nm.

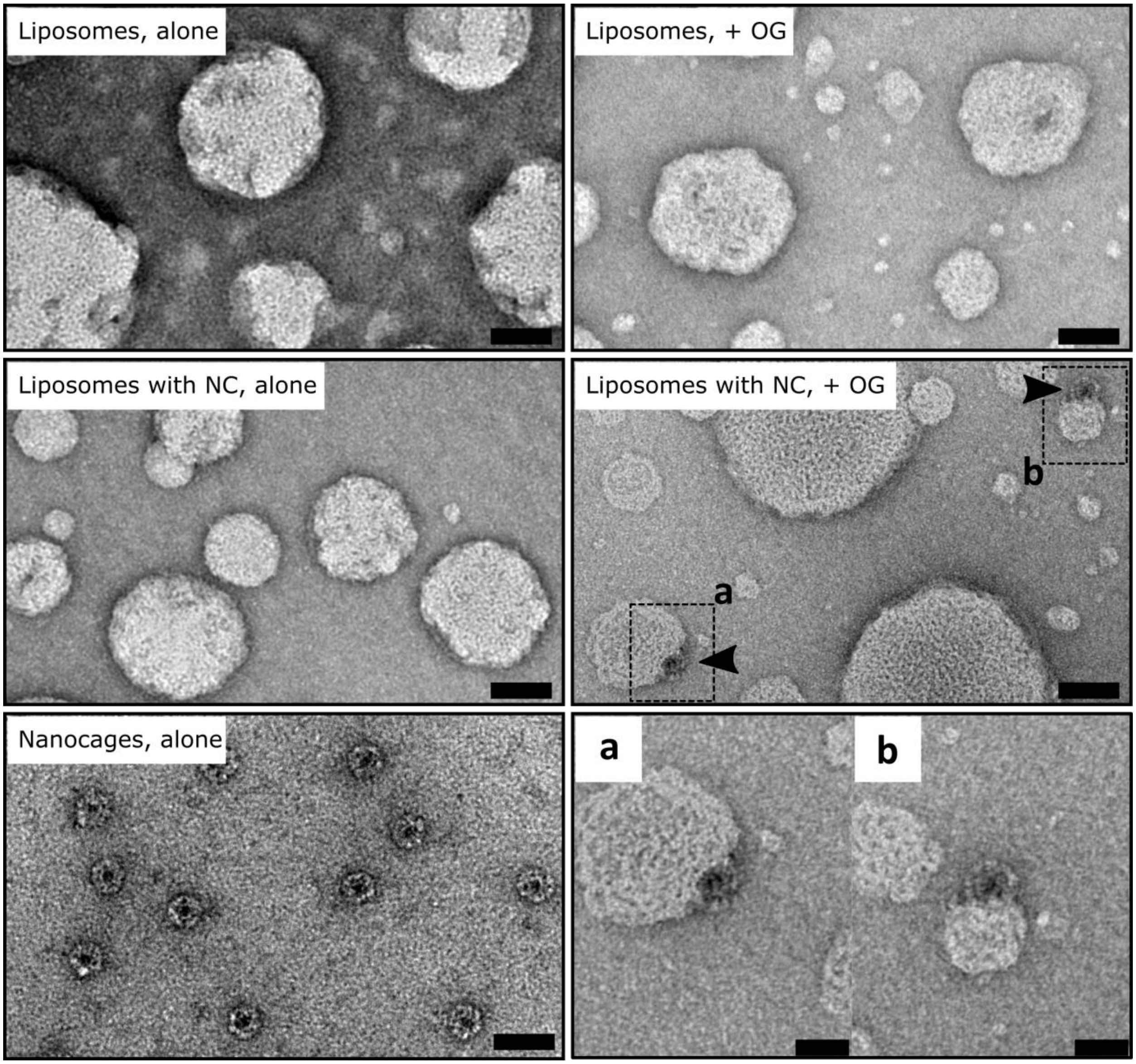

**Figure 9 | Nanocage release from liposomes.**

Transmission electron micrographs of empty, nanocage encapsulating liposomes and purified nanocages. OG (octyl-glucoside) was used to perforate liposomes in order to release internalized cargo. Lipid composition, POPC/POPS/Cholesterol. Scale bars, 50 nm. Zoomed in (a and b) scale bars, 25 nm.

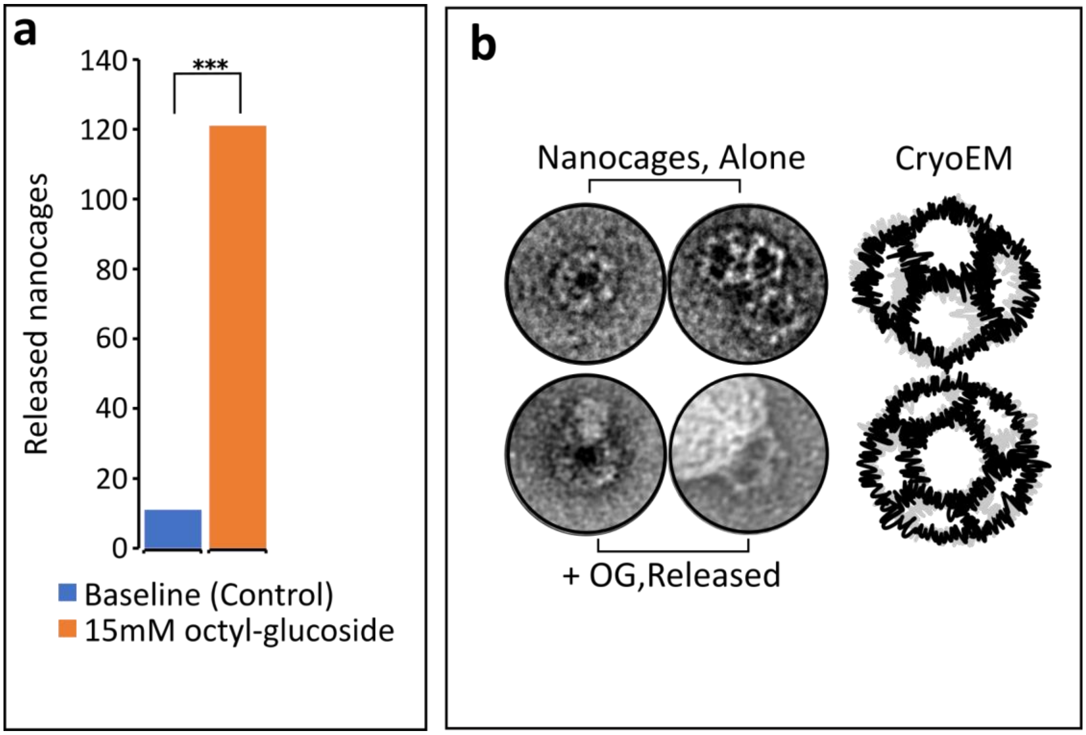

**Figure 10 | TEM investigation of nanocage encapsulation.**

**a**, Nanocage release assay as visualized by TEM and negative staining. Nanocages visible in the absence (blue), or presence (orange) of 15 mM octyl-glucoside. Total nanocage count represents 4 grid squares per condition; lipid composition: POPC/POPS/Cholesterol. *** $p < 0.001$. **b**, Images show nanocage structure before encapsulation (Nanocages, alone) and after encapsulation (OG, released). The structure of the encapsulated nanocages in this study matches the structure of the original design (based on 2D-class averages), as reported in[8]. Cartoon adapted from[8].

| | | | |
|---|---|---|---|
| Fluid Properties | $Density\ (\rho), \left[\frac{kg}{m^3}\right]$ | 1xPBS | $1007$ |
| | | Mineral Oil | $877$ |
| | $Viscosity\ (\mu), \left[\frac{kg}{m.s}\right]$ | 1xPBS | $89 * 10^{-5}$ |
| | | Mineral Oil | $0.0415$ |
| | $Interfacial\ Tension\ (\gamma), \left[\frac{kg}{s^2}\right]$ | Mineral Oil | $0.051$ |
| Channel Properties | $Geometry, [m]$ | Width | $250 * 10^{-6}$ |
| | | Height | $31 * 10^{-6}$ |
| Membrane Properties | $Pore\ Diameter, [m]$ | Membrane 1 | $100 * 10^{-9}$ |
| | | Membrane 2 | $5 * 10^{-6}$ |

**Table 1 | Model parameters for theoretical prediction of droplet size**
